# Non-motor effects of subthalamic nucleus stimulation in Parkinson’s patients

**DOI:** 10.1101/540328

**Authors:** Francesco Sammartino, Rachel Marsh, Ali Rezai, Vibhor Krishna

## Abstract

**Introduction:** The existing white matter connectivity analyses of the subthalamic region have mainly included the motor effects of deep brain stimulation.

We investigate white matter connectivity associated with the stimulation-induced non-motor acute clinical effects in three domains: mood changes, dizziness and sweating.

**Methods:** Using whole brain probabilistic tractography and seeding from the volumes of tissue activation, connectivity maps were generated and statistically compared across patients. The cortical voxels associated with each non-motor domain were compared with stimulation-induced motor improvements in a multivariate model. The resulting voxels maps were thresholded for false discovery (FDR q<0.05) and clustered using a multimodal atlas. To understand the role of local pathways in the subthalamic region, a group level parcellation was performed for each non-motor domain.

**Results:** The non-motor effects are rarely observed during stimulation titration: from 1100 acute clinical effects, mood change was observed in 14, dizziness in 23, and sweating in 20. Distinct cortical clusters were associated with each domain, notably mood change was associated with voxels in salience network and dizziness with voxels in visual association cortex. The subthalamic parcellation yielded a medio-lateral gradient with motor parcel being lateral and the non-motor parcels being medial. We also observed an antero-posterior organization in the medial non-motor clusters with mood changes (anterior), dizziness and sweating (posterior).

**Conclusion:** We interpret these findings based on the literature and foresee these to be useful for shaping the electrical field with the imminent use of steerable DBS electrodes.

## Introduction

Deep brain stimulation (DBS) of the subthalamic nucleus (STN) is efficacious for improving the motor symptoms of Parkinson disease. During DBS titration, clinicians adjust the stimulation to elicit acute clinical effects (ACE) which can either be efficacious or side effects. Generally, these ACE are in motor domains (e.g. improvement in tremor or motor contractions), however sometimes non-motor ACE are also observed. Although rare, several of these non-motor ACE have been previously reported, including episodes of mania, hilarity, and impulsive aggressive behavior [1]. These reported mood changes are both reproducible and reversible in a voltage-dependent manner [2]. While cortical connections associated with motor improvements following STN DBS have been extensively explored, the connectivity patterns underlying non-motor ACE remain largely unknown.

The topographical and functional organization of STN includes motor and non-motor subregions [3]. In vivo tractography indicated that similar anatomical subdivisions may exist in healthy humans [4]. Furthermore, STN DBS in PD patients was shown to modulate metabolic function in the motor, associative, and limbic cortical areas [5]. Although primate studies have also shown three distinct functional territories, recent 7T in-vivo studies have shown a degree of overlap between functional zones in the STN [6]. STN is a small nucleus and the electrical field from conventionally placed DBS electrodes likely spreads across functional territories. Further complicating matters is evidence that high-frequency stimulation may also activate white matter axonal tracts that extend outside the borders of the STN [7]. We hypothesized that unique cortical regions are associated with distinct non-motor ACE. We tested this hypothesis in a cohort of 24 PD patients with STN DBS using probabilistic tractography and simulated stimulation-associated volumes (VTA). Probabilistic tractography may be best suited to map the unique cortical clusters associated with non-motor ACE because of its ability of resolving multiple fibers orientations in voxels containing crossing-fibers.

## Materials and methods

### Study population

We acquired data from 24 patients with idiopathic Parkinson disease who underwent STN DBS at a single center (Center for Neuromodulation, The Ohio State University). These patients were deemed good surgical candidates following a multidisciplinary evaluation (including movement disorders neurology, neurosurgery, neuropsychology, and neuroradiology) and had at least 12 months follow-up after DBS insertion. Patients with surgery or device related complications were excluded. This study was performed in accordance with the Declaration of Helsinki. All patients gave informed written consent and this study was approved by the biomedical research institutional review board. We collected demographic and clinical information, and the levodopa equivalent daily dosage was calculated based on a previously reported formula [8].

From the reported profile of symptoms observed during DBS titration, we recorded each amplitude change separately for the following clinical domains: motor (rigidity, bradykinesia and tremor), mood change, dizziness (described as a sensation of spinning around and losing one’s balance while at rest) and sweating (accompanied by observable perspiration). The non-motor effects were patient-reported during stimulation adjustment while the motor changes were confirmed with clinical testing.

### Image acquisition

The pre-operative images were acquired on a Philips Achieva 3T scanner. Diffusion weighted images were acquired using an EPI sequence with inter-slice distance=0 mm; number of slices=71; voxel size=2.0 × 2.0 × 2.0 mm, TR=8100 ms; TE=68 ms; matrix size 128×128; 60 directions and 1 B0 image. High-resolution 3D anatomical scans were acquired using a T1 SENSE sequence (number of slices=170; voxel size 0.9×0.9×0.9 mm, TR=7492 ms; TE=3.6 ms; matrix size=240×240). A postoperative volumetric spiral CT scan without contrast (Siemens CT Somatom AS+, software version Syngo CT 2012B) was also acquired 4-6 weeks after electrode implantation (matrix size 512×512; voxel size 1×1×1; 200 images).

### Image pre-processing

Preprocessing was performed using FMRIB’s software library (FSL, http://www.fmrib.ox.ac.uk/fsl). After registering each diffusion volume to the b0 image using a 12 degree of freedom affine transform (FSL Eddy tool), the images were linearly aligned to each T1 in patient space and non-linearly registered to the MNI PD25 T1-MPRAGE template space using FNIRT. Similarly, the postoperative CT was also registered to this template space through the T1. The coordinates of the voxels corresponding to the individual contacts of the Medtronic 3389 DBS electrode (Medtronic, Inc, Minneapolis, MN) were selected [9]. Individual models of stimulation volume were created at each contact[10] using a Finite Element Model (FEM) by using specific amplitude and impedance for each patient.

### Analysis of cortical connections with probabilistic tractography

We performed probabilistic tractography in patient-space by seeding from each volume of tissue activation (VTA) mask (FSL Protrackx2 GPU version - n=25000 streamline fibers/voxel, curvature threshold=0.2, distance correction,3 fibers model, white matter inclusion mask) to the whole brain. We used a recently published cortical atlas to classify the tracks intersection with the cortex [11], and these masks were dilated to extend by 3 voxels into the white matter.

## Statistical analysis

We thresholded the raw tractography maps by the 60% of ‘robust threshold’ (FSL fslmaths) [12] and grouped each individual tractography map by the corresponding domain. For each pair of voltage changes (no ACE - ACE) we conducted a one-sample paired T-test using a non-parametric permutation-based approach with threshold-free cluster enhancement method (TFCE) as implemented in Randomise (http://www.fmrib.ox.ac.uk/fsl/randomise). We used 5000 permutations for statistical inference and adopted a corrected family-wise threshold of p<0.05 at the group level to produce the final thresholded voxel-wise maps grouped for each clinical domain.

A voxel-wise repeated measures ANOVA model using domain type as a categorical within-subjects variable and laterality as a categorical between-subject variable was run using AfNI 3dMVM [13] and the results were thresholded with FDR q<0.05 to yield the voxels significantly associated with each group. The final voxel maps were clustered using a multimodality cortical atlas [11].

### Creation of a group subthalamic nucleus map from intraoperative physiology data

The intraoperative microelectrode recordings data was analyzed for: location of STN (based on the appearance of STN cells and changes in background activity) and the motor STN sub-region (based on the presence of movement responsive or kinesthetic cells & tremor cells) and overlaid on patient MRI. We then registered the relative location of these cells on a validated template in MNI space from patients with PD (PD25 atlas) for comparison with the subthalamic area parcellation created from the previous analysis.

### Analysis of local connections with subthalamic area parcellation

The VTAs were averaged to create a group level mask in the subthalamic region. We created a group-dMRI template representing the cross-subject distribution of mean orientations from each subject’s Bayesian Estimation of Diffusion Parameters (BEDPOSTX, FSL suite). We masked the resulting maps for each domain using the T1-MPRAGE (1mm, MNI PD25 template) cortical ribbon (segmented with Freesurfer http://surfer.nmr.mgh.harvard.edu/), and used these voxels as target masks for connectivity driven parcellation of the group VTAs. The resulting connectivity matrix was clustered using k-means [14].

## Results

### Clinical outcomes

The demographic data and clinical outcomes are presented in Table 1. The UPDRS-III improved by 49.7% with a simultaneous decrease in levodopa equivalent by 54.8%.

**Table 1:**
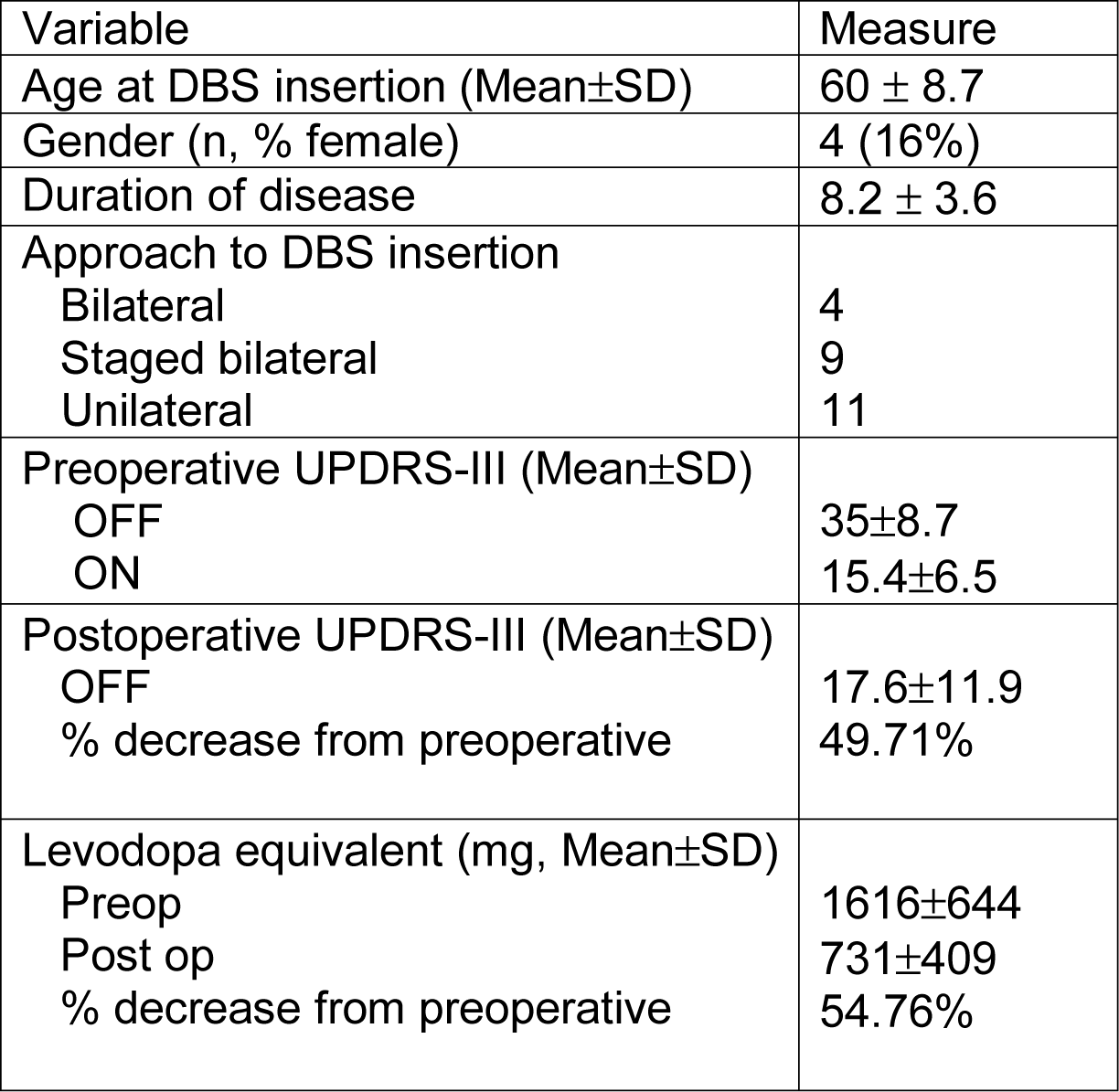
Demographics and clinical characteristics of the study population

### Stimulation data

From a total of 1100 ACE, 14 (1.3%) were classified as ‘mood change’ [observed in 5 patients], 23 (2.1%) ‘dizziness’ [observed in 8 patients] and 20 (1.8%) in the ‘sweating’ domains [observed in 7 patients] each. These were associated with 10, 18 & 11 unique voltage changes in the left hemisphere and 4, 5 & 9 voltage changes in the right hemisphere.

### Cortical anatomical connections associated with non-motor ACE

Probabilistic tractography from VTA voxels showed distinct patterns of cortical connectivity for each clinical domain.

The motor improvement domain (amplitude changes associated with improvement in rigidity, bradykinesia or tremor) was associated with voxels in the primary motor cortex (area 4) and superior frontal language area (area SFL), area 8, area 9 and area 47r anterior (Figure 1).

**Figure 1.**
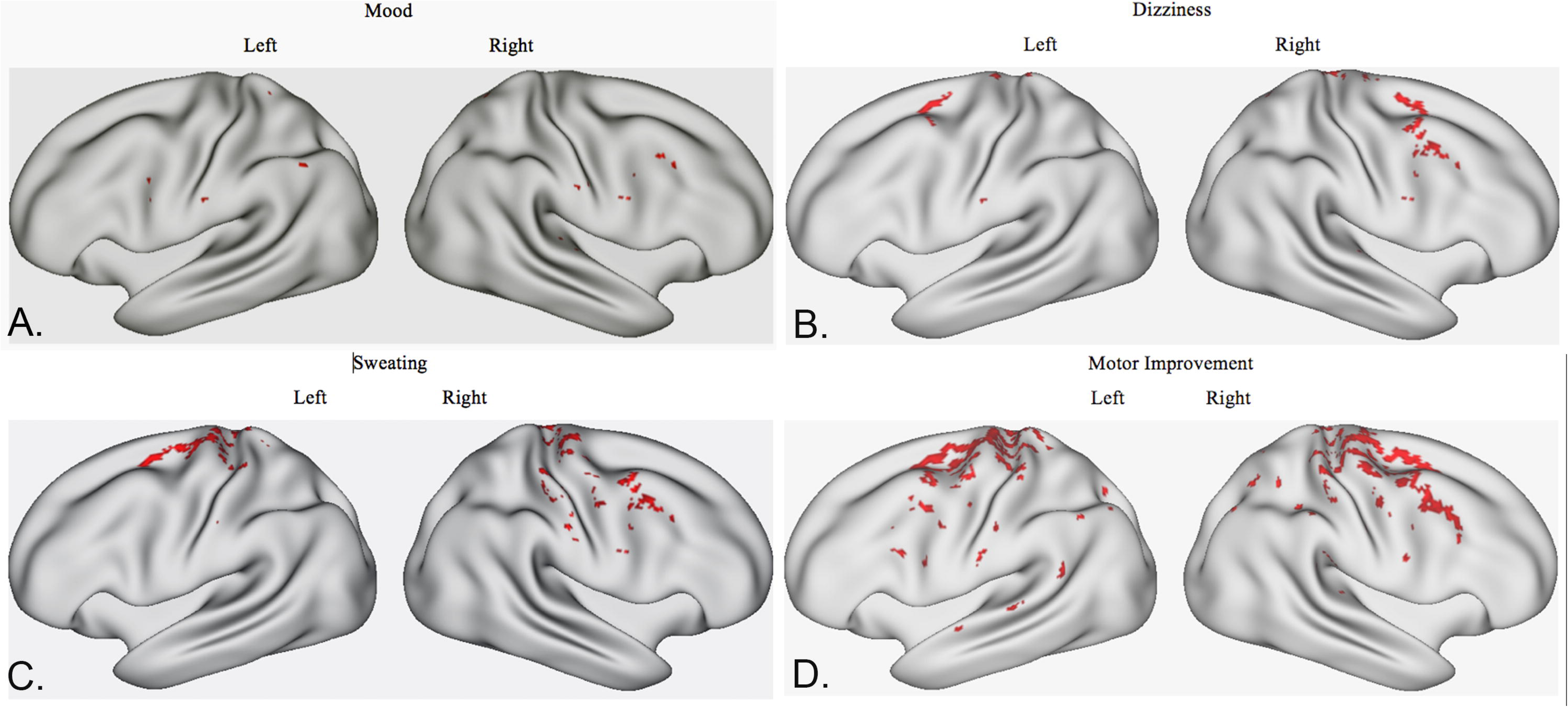
Cortical connections associated with non-motor ACE: Voxel maps for non-motor ACE in the mood (A), dizziness (B) and sweating (C) domains, compared with cortical voxels associated in motor improvement (D) after STN DBS.

In contrast, acute mood changes were associated with voxels in the insular (area Pol1 and Pol2) and temporal cortex (areas TE2 and TGd) (Figure 1).

Conversely settings changes associated with ‘dizziness’ were associated with voxels in Brodmann area 8, temporal cortex (areas TE2, TGd), Brodmann area 9 and insula (area Pol2) (Figure 1).

Finally, sweating domain was associated with voxels in temporal cortex (areas TE2 and TG dorsal), primary motor cortex (area 4) and areas 8 – 9 (Figure 1). The cortical areas results are also ordered in Table 2.

**Table 2:**
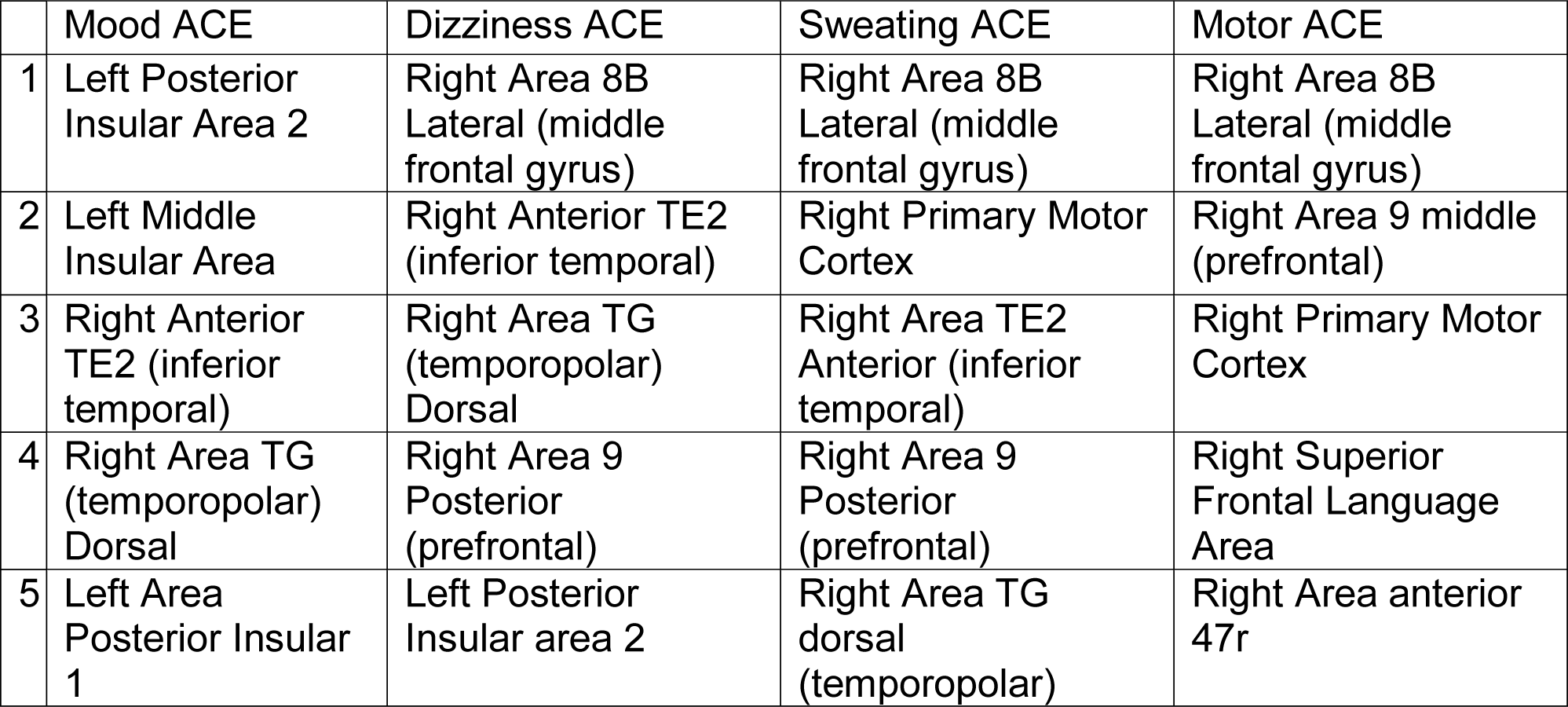
Cortical areas with highest white matter connections in non-motor and motor domains

### Local anatomical connections associated with non-motor ACE

From the parcellation of the group VTA masks, clusters associated with improvement in motor symptoms (rigidity, bradykinesia, tremor) after DBS were localized in the lateral portion of the mask [MNI coordinates of the center of gravity: X=−13.7(13.7) - Y=−13.5(−13.5) - Z=−8.7(−8.7)] (Figure 2 A-B).

**Figure 2.**
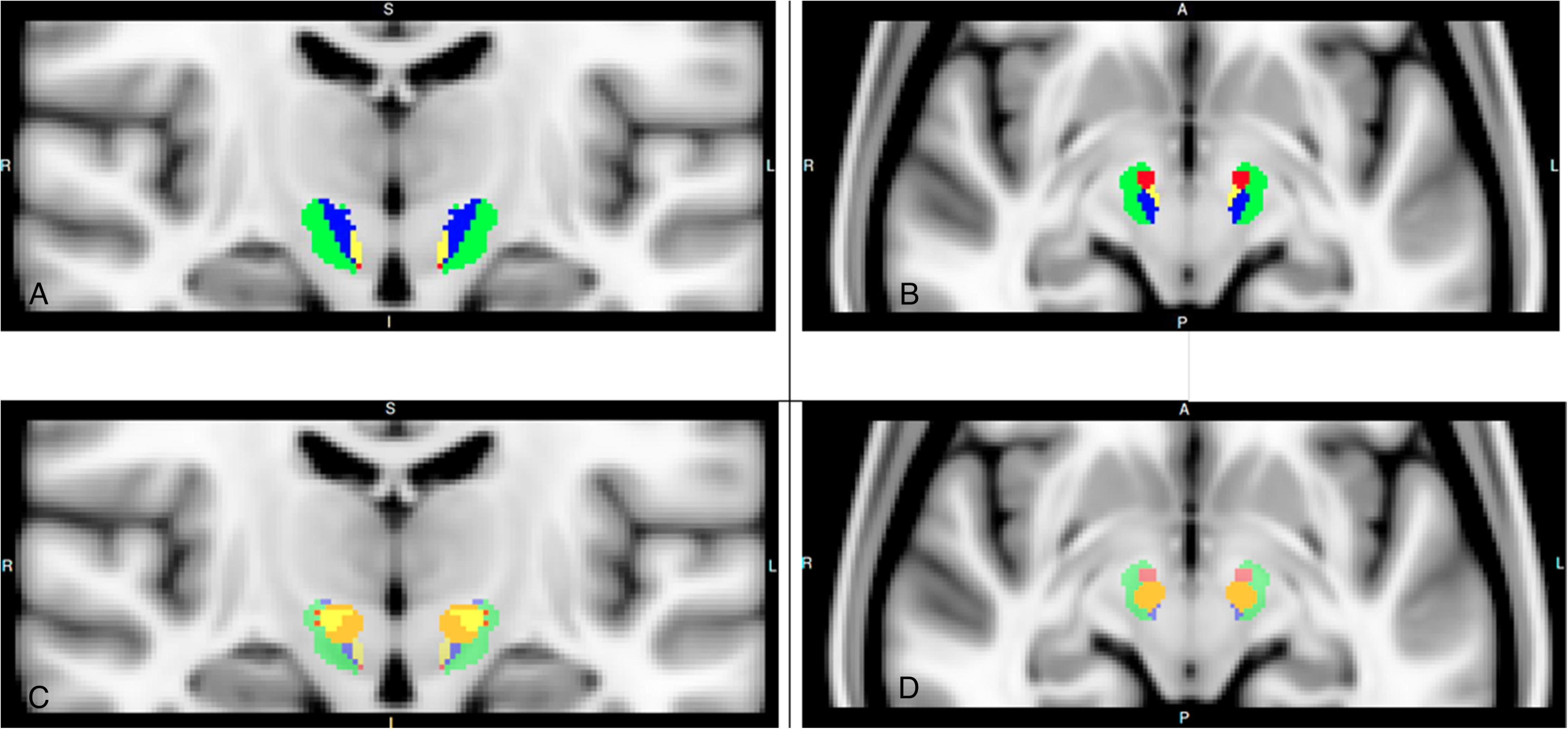
Local pathways associated with non-motor ACE: Group level parcellation of the volumes of tissue activation for side effects in mood(red), dizziness(yellow), sweating (blue) and motor improvement (green) (A-B), compared to the group level STN map defined by neurophysiology data (orange color, within the motor territory) (C-D).

Conversely clusters with connectivity to the cortical areas associated with non-motor ACE were located more medially. Voxels associated with mood changes were clustered more anterior and medially [MNI coordinates of the center of gravity: X=−11.7(11.7) - Y=−10.5(−10.5) - Z=−8.4(−8.4)]. The cluster associated with dizziness was located dorsomedially [MNI coordinates of the center of gravity: X=−10.9(10.9) - Y=−11.7(−11.7) - Z=−5.2(−5.2)] while the cluster associated with sweating was located posterior and ventrally [MNI coordinates of the center of gravity: X=−12.3(12.3) - Y=−14.7(−14.7) - Z=−6.1(−6.1)].

There was no statistically significant difference in the volume of the clusters between the two hemispheres (paired t-test, two tailed, p>.05). These non-motor parcels were extending outside the boundaries of the group STN map (C - D).

## Discussion

Here we investigate the connectivity substrates of non-motor side effects observed during stimulation titration in PD patients with STN DBS. Using an unrestricted whole brain connectome built using probabilistic tractography, we observed unique cortical clusters associated with mood, sweating and dizziness domains. We then identified discrete clusters in the subthalamic region uniquely associated with cortical clusters in each domain. We believe that these findings expand current knowledge, providing further insights into shaping the stimulation field through steerable DBS electrodes for broadening the therapeutic window.

### The cortical and subcortical connections of STN

The STN balances inhibitory and excitatory signals to modulate basal ganglia-thalamocortical outflow and control complex behaviors. GABAergic neurons from the globus pallidus pars externa (GPe) contribute major inhibitory afferent connections to the STN, as motor and mood-related portions of the GPe innervate corresponding counterparts in the dorsolateral and ventromedial portions of the STN, respectively [15]. Pallidal GABAergic afferents to the STN initiate IPSPs through the activation of post-synaptic GABA_A_ receptors while presynaptic effects are mediated through GABA_B_ receptors. These GABAergic afferents are balanced by glutamatergic input from the cerebral cortex and thalamus. Primate studies indicate that the cortical afferents arise primarily from the primary motor cortex, supplementary motor area (SMA), and premotor cortices and project to the dorso-lateral motor region of the STN. In rodents, cortical afferents from the prefrontal cortex and prelimbic-medial orbital areas also innervate the medial STN. In primates, it is unclear whether these afferents are direct but their density of cortical terminals in non-motor STN matches the density seen in motor STN in primates [16]. Furthermore, primate studies have demonstrated that thalamic projections from the parafascicular and centromedian nuclei project topographically to the medial rostral and dorsolateral motor STN territories, respectively. The centromedian (CM) and parafasciular nuclei (Pf) of thalamus also provide excitatory input, contacting distal dendrites of the STN. Despite the apparent anatomical segregation of cortical afferents to the STN, the nucleus is small and contains neurons with large dendritic trees, rendering it a synaptically convergent system between different functional subregions. Recent multimodal investigations with the integration of neurophysiology and neuroimaging (both DTI and fMRI) also indicate the robust connectivity of STN and its neighboring white matter to both sensorimotor areas and regions involved in cognitive and emotional/behavioral functions like the prefrontal cortex and the SFL area [10]. This diffuse connectivity led us to perform a whole brain tractography analysis without restricting the target masks to specific cortical areas.

### Mood-related ACE may be mediated by the connections to the salience network

In addition to the motor improvements after STN DBS, several groups have reported stimulation-related changes to baseline mood [17] and cognitive performance. It should be emphasized that the altered susceptibility to mood changes in PD may also be related to a variety of other factors including improved motor disability and medication reduction [18]. However, the mood and behavioral effects STN DBS are well known and therapeutic trials have investigated its application for behavioral disorders like OCD [19]. The mechanisms however remain unclear. Previous work postulates that inadvertent co-activation of axonal tracts in the medial forebrain bundle (MFB) mediated by STN stimulation may drive DBS-induced acute hypomania in PD patients. Dopaminergic neurons within the MFB that contribute to mesolimbic and mesocortical circuitry originate in the ventral tegmental area, located anterior and medial to the STN. For ACE in mood domain, we observed cortical clusters in the insular cortex and salience network [20]. Clusters were identified in posterior insular areas Pol1 and Pol2, as well as middle insular area (MI). The insular cortex is a critical hub in the salience-processing network, detecting behaviorally relevant stimuli from ascending interoceptive and autonomic afferents to facilitate an appropriate response. Aberrant insular activation is associated with inappropriate salience detection and disrupted attentional responses that underlie neuropsychiatric disorders. The insular cortex can be segregated into distinct subdivisions based on connectivity patterns, further elucidating its functional contribution to many diverse processes in salience processing. The MI is localized within the anterior insular cortex (AIC) and functional studies have implicated it in tracking emotion and perceptions through the incorporation of conscious thought and autonomic nervous signals [21]. The AIC demonstrates extensive connections to the amygdala, anterior cingulate cortex (ACC), orbitofrontal cortex (OFC), and temporal lobes. The anterior ventral insula and ACC uniquely also contain the Von Economo neurons (VENs). These neurons are believed to relay efferent projections from the AIC and ACC to frontal and temporal regions, thus facilitating rapid cognitive-emotional responses. The preferential degeneration of VENs in frontotemporal dementia further indicates their involvement in awareness, emotional response, and self-control [22]. Our analysis also indicates a role for Pol1 and Pol2, overlapping regions in the posterior insular cortex (PIC). Unlike the AIC, which drives perception, the PIC determines stimulus intensity (e.g. pain) in salience processing. This is likely mediated by somatosensory activation, explained by significant connections with spinothalamic fibers via primary and secondary somatosensory cortical areas. Our results favor steering the electrical field away from the most ventral portion of the subthalamic region to avoid evoking unpleasant mood changes.

### Dizziness may involve STN connection to the visual processing network

Previous reports have documented dizziness during intraoperative STN stimulation mainly in medial electrode locations adjacent to the sites inducing eye deviation [23]. We identified cortical clusters in the temporal polar cortex (TPC), TE2a (inferior temporal sulcus) and TGd (dorsal) associated with acute dizziness during acute stimulation adjustment. The posterior temporal cortex contains a hierarchy of visual association areas primarily extending anteriorly from the occipital cortex. Neurons along the occipitotemporal cortical pathway in the inferior temporal area (TE) respond selectively to object identification, but are unresponsive to spatial aspects of stimuli [24].

Similarly, lesions of the inferior temporal cortex cause deficits in visual discrimination tasks but not visuospatial tasks [25]. Here we conjecture that acute dizziness may be mediated through disruptions in the ventral stream clusters in the inferior temporal cortex, associated with the visual processing network that enables conscious visual perception of self-motion. Another possibility could be underlying vestibular dysfunction coupled with an increased visual dependence causing visual vertigo. It is well known that patients with PD exhibit reduced vestibular responses compared with controls [26] and in situations with excessive visual motion, may have a limited ability to fully compensate for the maladaptive vestibular response, presumably inducing a sensation of dizziness.

### Sweating ACE may be mediated by the STN connections to temporal cortex and hypothalamus

Autonomic dysfunction with sweating is common in patients with Parkinson disease and it is not improved by DBS e.g. among 77 PD patients 64% reported sweating disturbances, either hypohidrosis or hyperhidrosis and often asymmetric, occurring predominantly in *off* periods and in *on* periods with dyskinesias [27]. It is not entirely clear whether it is a part of the disease process or medication side effects. It may be mediated peripherally by postganglionic fibers in patients with long-standing disease. However, it is also possible that it may have a more central mechanism. Sweating has previously been reported in response to stimulation in a PD patient [28]. We found that cortical areas TE2a and TGd were correlated with acute sweating side effects, together with the postero-medial subthalamic parcel. The postero-medial STN is adjacent to the lateral hypothalamic area, which contains the orexinergic nucleus mediating thermoregulation. The lateral spinothalamic tract is the major thermal and pain sensitivity afferent pathway projecting to the hypothalamic thermoregulatory centers. Thus, stimulation of contacts in the postero-medial STN may lead to DBS-mediated activation of autonomic changes via spread to the hypothalamus. Similarly, intraoperative stimulation in the medial area of the STN induced vegetative side effects (nausea, heat sensation, sweating), also exhibited to a lesser extent following stimulation of the STN [29]. It is also known that neurons in the temporal cortex can modulate the central autonomic network. Serial measurements of palmar galvanic skin reflex [30], both in non-human primates than in humans, have also revealed the premotor and temporal cortex together with the anterior portion of the hypothalamus/prechiasmatic region are involved in sweating. The exact mechanism of stimulation-induced sweating remains unclear.

#### Limitations

This is a retrospective tractography study to investigate whether non-motor ACE observed during STN DBS titration are associated with connections to unique cortical regions. However, we are unable to conclude whether changes in activity of these cortical clusters underlies the observed clinical effects. Such investigations will require functional studies with functional MRI or magnetoencephalography. It is also unclear whether ACE predicts chronic stimulation outcomes which are often complicated by factors like habituation to chronic stimulation. These findings will need prospective validation in a larger cohort to understand their significance for DBS targeting and shaping of stimulation field.

## Conclusion

Using probabilistic tractography and modeling of stimulation volumes associated with STN DBS, we observed distinct clusters both cortically and locally in the subthalamic area associated with non-motor ACE associated with specific voltage changes in a cohort of PD patients. Though with important limitations, these findings may help understand the mechanisms of common non-motor effect of DBS and ultimately also help avoid these side effects with automated algorithms for an efficient shaping of the stimulation volume.

## Acknowledgments

We thank Qinwan Rabbani and Thomas Geist for help with data collection.

## Funding sources

1. Discovery themes initiative funding, The Ohio State University
2. Neurological Research Institute, The Ohio State University

## Authors’ Roles

1. Research project: A. Conception, B. Organization, C. Execution; Francesco Sammartino, Vibhor Krishna
2. Statistical Analysis: A. Design, B. Execution, C. Review and Critique; Francesco Sammartino, Vibhor Krishna
3. Manuscript: A. Writing of the first draft, B. Review and Critique. Francesco Sammartino, Vibhor Krishna, Rachel Marsh, Ali Rezai

## Abbreviations

ACE: acute clinical effects
VTA: volume of tissue activation
STN: subthalamic nucleus
DBS: deep brain stimulation

## Notes

Financial Disclosure/Conflict of Interest concerning the research related to the manuscript: the authors do not have disclosures or conflicts to declare.

